# Bayesian inference of cancer driver genes using signatures of positive selection

**DOI:** 10.1101/059360

**Authors:** Luis Zapata, Hana Susak, Oliver Drechsel, Marc R. Friedländer, Xavier Estivill, Stephan Ossowski

## Abstract

Tumors are composed of an evolving population of cells subjected to tissue-specific selection, which fuels tumor heterogeneity and ultimately complicates cancer driver gene identification. Here, we integrate cancer cell fraction, population recurrence, and functional impact of somatic mutations as signatures of selection into a Bayesian inference model for driver prediction. In an in-depth benchmark, we demonstrate that our model, cDriver, outperforms competing methods when analyzing solid tumors, hematological malignancies, and pan-cancer datasets. Applying cDriver to exome sequencing data of 21 cancer types from 6,870 individuals revealed 98 unreported tumor type-driver gene connections. These novel connections are highly enriched for chromatin-modifying proteins, hinting at a universal role of chromatin regulation in cancer etiology. Although infrequently mutated as single genes, we show that chromatin modifiers are altered in a large fraction of cancer patients. In summary, we demonstrate that integration of evolutionary signatures is key for identifying mutational driver genes, thereby facilitating the discovery of novel therapeutic targets for cancer treatment.

---

Since the 1970s, tumors have been considered the product of evolutionary forces such as positive selection of highly proliferative cancer genotypes or negative selection of nonadaptive cancer genotypes^1^. Analogous to the evolution of multi-cellular organisms, random somatic mutations in cancer cells interplay with natural selection, creating phenotypic diversity and allowing for adaptation^2,3^. It has been shown that this process of clonal evolution follows different paths depending on the background genotype of patients^4,5^, the tissue microenvironment^6^, and the functional redundancy of acquired somatic mutations^7^. This leads to increased molecular diversity, ultimately contributing to intra- and inter- tumor heterogeneity^3^. This heterogeneity, ubiquitously present in tumor types^8,9,10,11^, hampers the identification of driver genes and hence limits the number of therapeutic targets to be detected^12^.

Next generation sequencing (NGS) allows mutational screening across thousands of tumors uncovering the extent of cancer heterogeneity^13,14,15^. Recent methods have used NGS to infer tumor phylogenies by estimating the cancer cell fraction (CCF) of variants present only in the tumor (somatic mutations)^16,17,18,19,20^. Consequently, evaluation of solid tumors has uncovered common mutations coexisting with region-specific mutations^21,9,22,15^, and studies in hematological malignancies have revealed clonal and sub-clonal variants in the same sample^8,23,24^. These efforts have shed light into the extent of sub-clonal versus clonal genetic variation observed across tumors, highlighting that sub-clonal mutations accumulate predominantly in a neutral fashion^52^ and that the average cancer cell fraction (CCF) is higher for driver than for passenger mutations^25^. Nonetheless, CCF has not been applied as a feature for the identification of mutational driver genes.

Current solutions for identifying driver genes rely on the recurrent mutation of genes across a large number of cancer patients^26^, the genomic context where they occur^13^, the functional impact of mutations^27^, and the clustering of mutations within functional protein domains^28^. However, statistical methods based on mutation recurrence and context alone have not been able to classify infrequently mutated genes as drivers^3^. To this end, methods based on molecular selection signatures, such as functional impact and mutation clustering, have been combined to identify these elusive driver genes^29^, but without considering CCF. A large number of tumor samples will continue to be sequenced at increasing depth of coverage, allowing for accurate identification of sub-clonal mutations and, therefore, requiring integrative models to differentiate early and late driver from passenger genes. Knowledge of the driver gene landscape is key to improve diagnosis, selection of treatment, monitoring of progression, and identification of treatment resistant sub-clones at earlier time points^30^.

Here, we present cDriver, a novel Bayesian inference approach to identify and rank mutational driver genes using multiple measures of positive selection. We benchmark our results against standard tools on public tumor datasets. Finally, we apply cDriver to 6,870 cancer exomes to uncover associations between driver genes and tumor types, identifying novel connections highly enriched for chromatin modifying proteins, expanding the current set of prognostic markers for cancer treatment.

## Results

### Evolutionary signatures used by cDriver

To identify driver genes, we have developed cDriver (**Supplementary Fig. 1, online methods**), a Bayesian model that integrates signatures of selection of somatic point mutations (SNVs and short indels) at three levels: i) population level, the proportion of affected individuals (recurrence), ii) cellular level, the fraction of cancer cells harboring a somatic mutation (CCF), and iii) molecular level, the functional impact of the variant allele (Fig. 1). cDriver is the first method that incorporates recurrence (Fig. 1a), CCF (Fig. 1b), and functional impact (Fig. 1c) to identify mutational driver genes based on a probabilistic framework. By definition, a driver event involves the acquisition of a somatic mutation conferring a selective advantage at the cellular level^2^, therefore mutations found at the root of the tumor evolutionary tree, or mutations leading to a selective sweep, will consistently be at high CCF. Conversely, it is possible that passenger somatic mutations in the last common ancestor of the cancer clone (i.e. predating malignant transformation) hitch-hiked with a driver event^31^, reducing discriminating power of CCF as a signature of positive selection.

**Figure 1.**
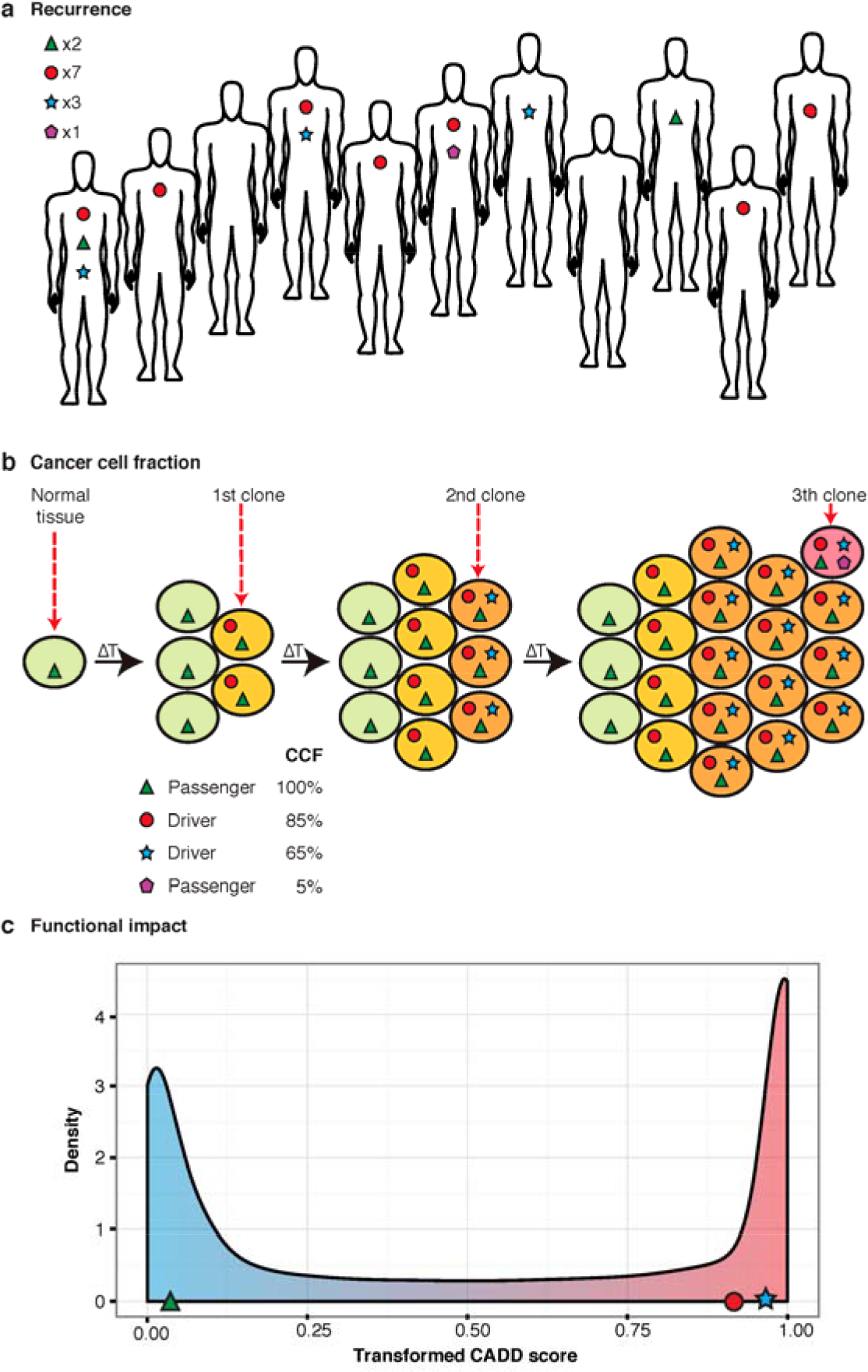
Signatures of positive selection observed from tumor sequencing data. (**a**) Large-scale sequencing experiments of patient cohorts reveal the mutational landscape of a cancer across a population. Somatic mutations under positive selection (circle and star) are expected to be more frequent than somatic mutations that confer no selective advantage (triangle and pentagon). As a result, most of the current algorithms consider recurrently mutated genes as drivers and randomly mutated genes as passengers. (**b**) Illustrative model of clonal evolution showing four time points. Each clone is represented by a unique genotype, and is depicted as a group of cells (ellipsoids) with the same background color. Shapes inside the cell represent mutations. Two types of mutations under positive selection are illustrated: a tumor-initiating driver (red circle) and a late-driver causing clonal expansion (blue star). The initial driver mutation causes the emergence of the first malignant clone (last *onko*-common ancestor, LOCA) and it propagates to all daughter cells, thus having a high cancer cell fraction (CCF) at all time points. The second driver mutation confers a selective advantage over the rest of the clones, generating a selective sweep in the last time point. Two types of passenger mutations are shown: early passengers or hitchhikers (green triangle) present at a high CCF since they appeared before the emergence of the LOCA and late passenger (purple pentagon) present only in a small fraction of cancer cells. The CCF value describes the total fraction for each mutation at the last time point. (**c**) Highly damaging mutations are expected to be under selection given they disrupt the normal protein function. In contrast, passenger mutations are mostly neutral and are not expected to have a bias towards high functional damage. In this study we integrate signals depicted in a-c in one model for driver gene identification.

To test if CCF discriminates between driver and passenger mutations and is not solely a signature of timing, we compared the CCF distribution of somatic mutations in a set of published driver and non-driver genes (Fig. 2) (**Online methods**). We observed that the median CCF of nonsilent mutations was significantly higher in driver genes than in non-driver genes in a pooled set of 16 TCGA tumor studies (Mann-Whitney P Value < 1e-323), in curated data of 12 tumor types^32^ (P Value = 1.5e-120), in a solid tumor (BRCA, P Value = 1.7e-86), and in a hematological malignancy (CLL, P Value = 4.7e-07). Importantly, a significant difference was also observed when comparing the median CCF of nonsilent to silent mutations in driver genes, demonstrating that positive selection is specifically acting on nonsilent mutations in driver genes. Moreover, there is no difference in median CCF between nonsilent mutations in passenger genes and silent mutations in any gene, indicating that the observation is not caused by a systematic bias (e.g. gene length bias). We observed similarly significant results across 14 out of 16 tested tumor types (**Supplementary Fig. 10**). Considering that a fraction of passenger mutations likely occur prior to malignant transformation, CCF alone may be insufficient to pinpoint all driver genes. Similarly, not all oncogenic mutations show high functional impact scores (**Supplementary Fig. 13**) and not all driver genes are highly recurrent in a cohort. cDriver overcomes these limitations by integrating recurrence, CCF and functional impact as signatures of positive selection at a population, cellular, and molecular level, respectively (**Online methods**).

**Figure 2.**
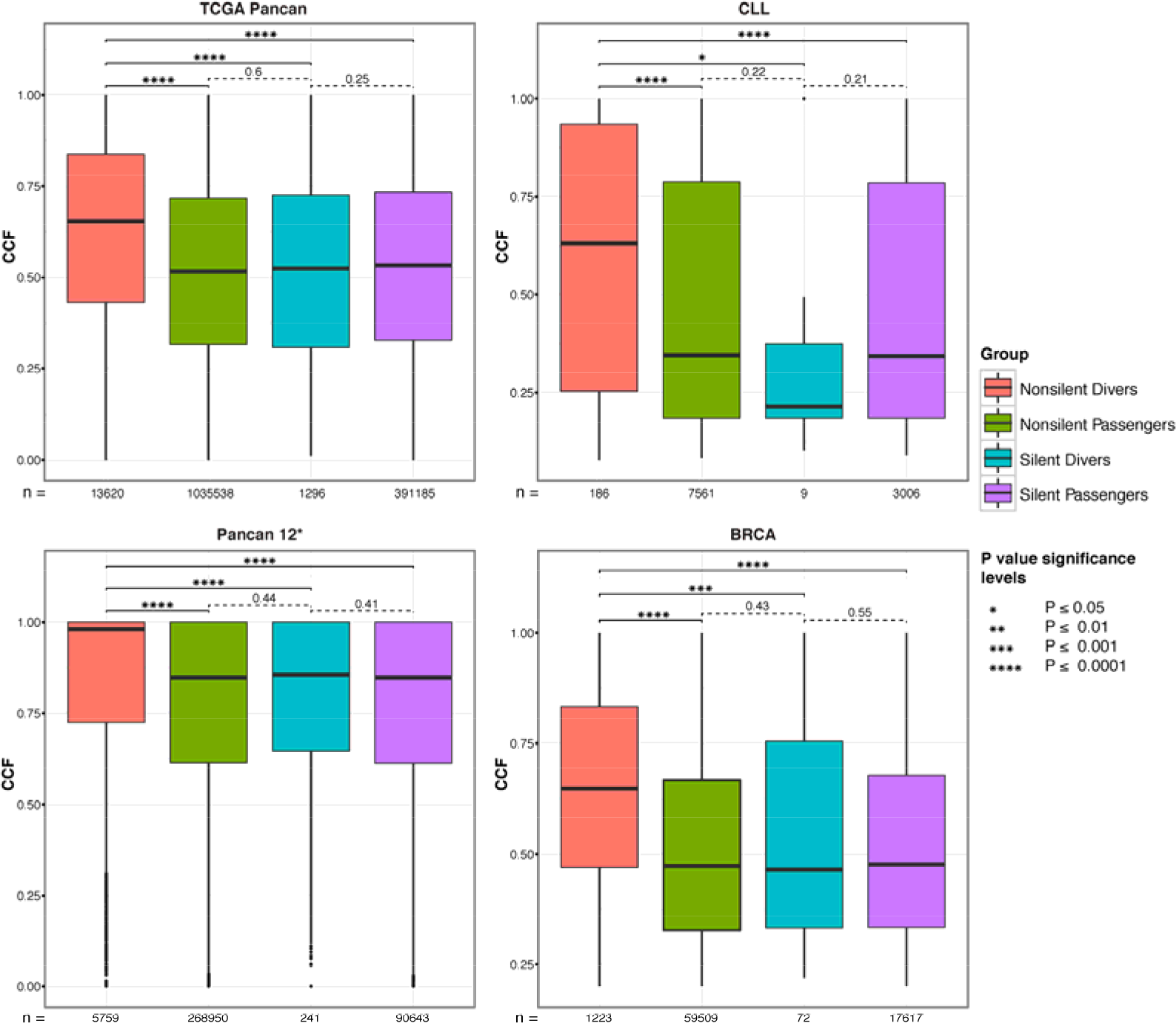
CCF distribution for four groups of somatic mutations in four cancer datasets. We obtained the CCF distribution for nonsilent driver, nonsilent passenger, silent driver, and nonsilent passenger gene mutations and compared the significance of the differences between each pair of them. CCF of nonsilent driver mutations is significantly higher compared to all other groups. Importantly, CCF of nonsilent driver is significantly higher compared to silent mutations in driver genes and the latter were not significantly different from silent or nonsilent mutations in passenger genes (*Pancan12 represent the highly filtered dataset published in^32^).

### Benchmarking cDriver performance in different tumor types

To assess the performance of cDriver, we benchmarked against four frequently used driver gene identification methods using precision, recall, and F-score (**Online methods**). cDriver, OncodriveFM^27^, OncodriveCLUST^28^, MuSiC^26^, and MutsigCV^13^ were run on three public cancer datasets. These datasets consisted of 762 cases of breast cancer (BRCA)^32^, 385 cases of chronic lymphocytic leukemia (CLL)^33^, and 3,205 cases from a pool of 12 tumor types (Pancan12)^32^ (**Supplementary table 1**). Results of each method for each dataset were sorted by P values or posterior probabilities to compare ranked driver genes.

### Benchmarking in breast cancer (BRCA) and chronic lymphocytic leukemia (CLL)

To benchmark competing methods when analyzing solid and hematological tumor data, we assembled a list of 33 and 22 gold standard (GS) genes for Breast Cancer (BRCA) and chronic lymphocytic leukemia (CLL), respectively (**Supplementary Table 2, Online Methods**). We found that cDriver outperforms all other methods in both BRCA (Fig. 2a) and CLL (Fig. 2b), showing the highest F-score, as well as similar or better recall and precision as the second best method (**Supplementary Fig. 2**).

To reveal if cDriver was able to identify significant mutational driver genes not highly ranked by other methods, we developed a model to estimate a rank cutoff for a desired FDR (**Online methods**). At significance level (FDR 0.1), we found 36 driver genes for BRCA and 23 for CLL. We next looked at genes in the GS (**Fig 3c, d, upper panel**) and cDriver significant genes not present in the GS (**Fig 3c, d, lower panel**). On one hand, we observed that five GS BRCA genes and six GS CLL genes were not significant in any method, suggesting that these genes are likely affected by variation other than somatic point mutations. On the other hand, cDriver significantly found 12 genes in BRCA and 11 genes in CLL not present in the tumor-specific GS (**Fig. 3c, d, lower panel**). Although most of these genes were highly ranked by at least one other method, two genes were highly ranked and significant only by cDriver, *KDM6A* and *EGR1*. The former was recently reported to play a role in a rare aggressive breast cancer^34^, while the latter was found related to CLL using gene expression and network analysis^35^, reaffirming their putative role in tumor etiology. Several of the remaining 21 genes not in the GS, such as *MYH14, MED23*, and *ZFP36L1* in BRCA, and *ZNF292, FUBP1*, and *DTX1* in CLL, had also been implicated in tumor development^36,33,37,38^.

**Figure 3.**
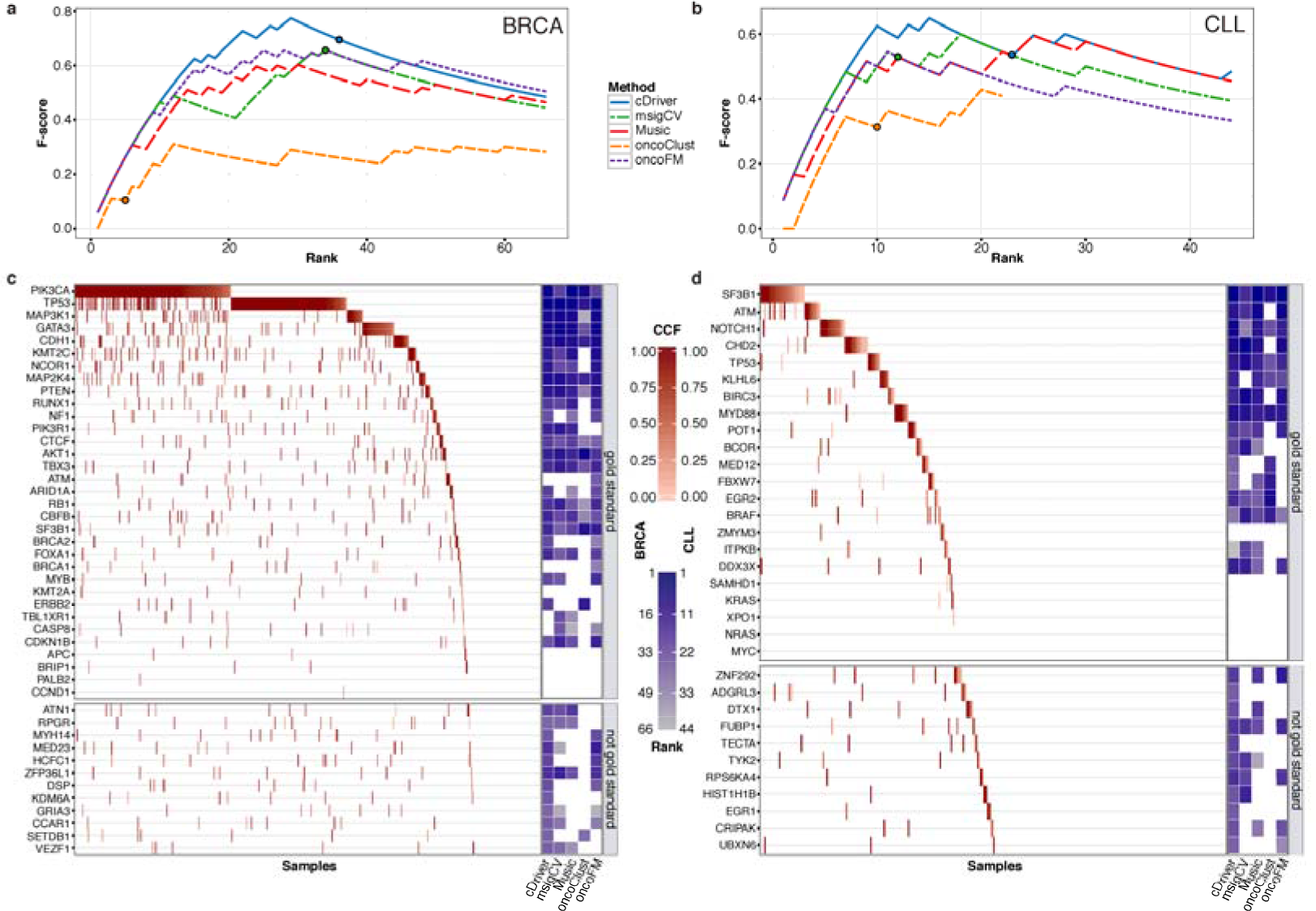
Benchmarking of cDriver and other driver identification methods in breast cancer (BRCA) and chronic lymphocytic leukemia (CLL) datasets. F-score for cDriver (solid blue line) and four other driver identification algorithms using BRCA (**a**) and CLL (**b**) datasets. Results of each method were transformed to ranks by ordering P values or posterior probabilities. The P value cutoff for significance is shown as a circle in each of the curves. For visualization, F-score is shown to rank 66 for BRCA and 44 for CLL (twice the number of genes in the gold standards), since all methods reach the F-score peak before these ranks (c, d). We compared the results for all methods irrespective of the P value using only the ranking for BRCA (c) and CLL (d). Gold standard genes were ordered by mutation frequency and samples were ordered by cancer cell fraction (CCF). The CCF of each mutation in each gene-patient pair is indicated by the red color gradient. On the right, gene rankings of each algorithm are indicated by the blue color gradient. White means that this gene was not ranked under 66 for BRCA (**c**) and 44 for CLL (**d**). At the bottom of figures c and d results for genes not present in the gold standard but highly ranked by cDriver are shown.

### Benchmarking in a pan-cancer dataset of 12 tumor types

Capitalizing on the large number of whole exome sequencing (WES) studies published by TCGA, we benchmarked all methods on a high quality (curated) dataset of 12 tumor types (Pancan12)^32^ using five published gold standards (GS)^39,32,40,30,28^ (**Online methods**). Across ˜3,200 samples, cDriver and oncodriveFM performed best in F-score using Cancer Gene Census (Fig. 4a) with and without filtration of non-expressed genes (**Supplementary Fig. 3**). Noteworthy, only MuSiC benefited extensively from this post-filtration step. In addition, cDriver outperformed all other methods amongst the top 50 ranked genes across all gold standards using F-score, recall, and precision (**Supplementary Fig 4**). We also noticed that significance thresholds used by different methods often do not coincide with their respective F-score peak independently of the GS used, e.g. Music and OncodriveFM are often far from optimal F-score at significance level (Fig 4a **circles in F-score curves, Supplementary Table 3**). At significance level cDriver suggests a cutoff at 418 genes for Pancan12 (**Supplementary Fig. 11**) close to the optimal F-score. At significance level cDriver had the best F-score in three out of five GS and at least the second-best F-score in all GS (**Supplementary Table 3**).

**Figure 4.**
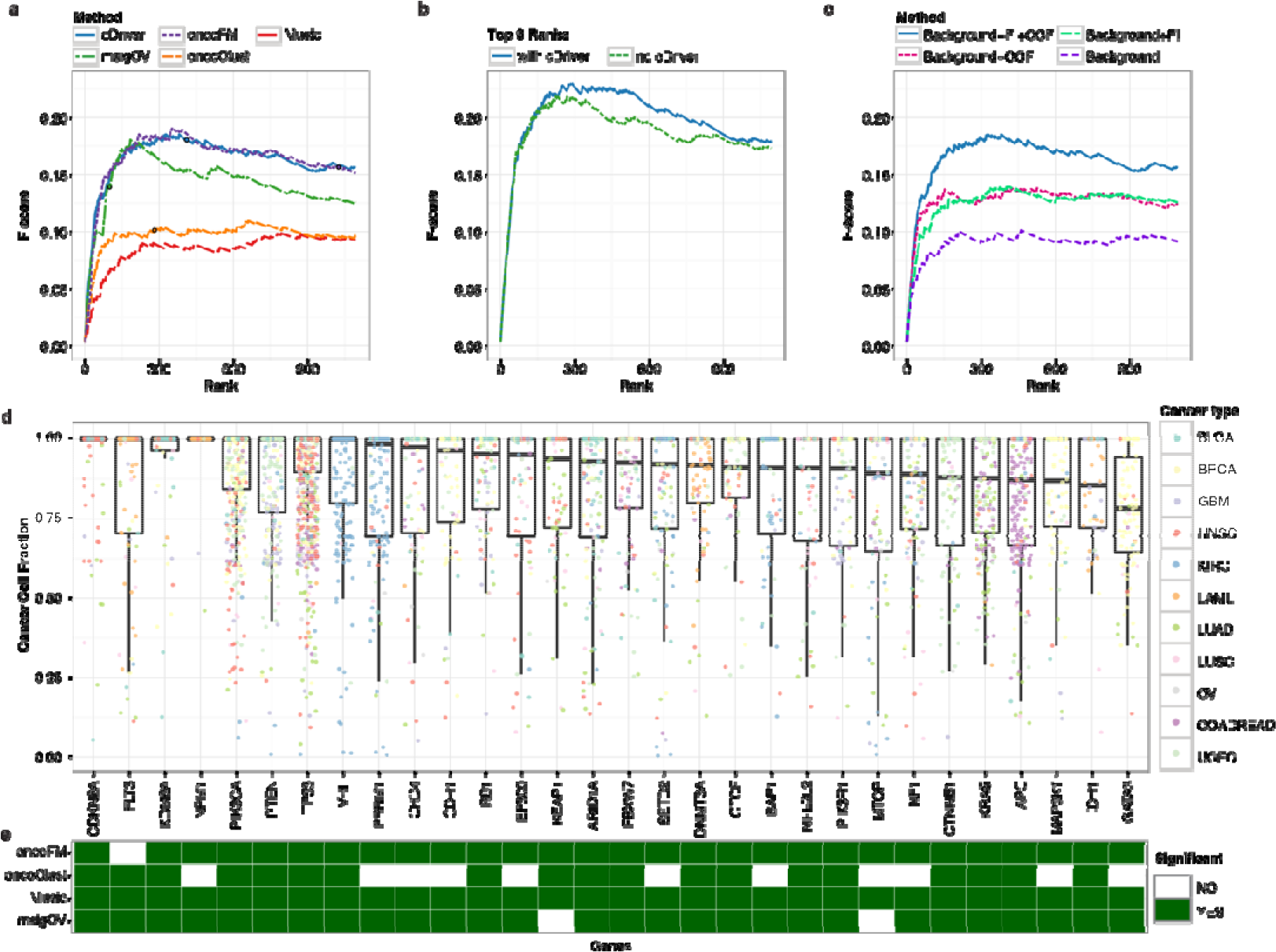
cDriver results and comparison with other methods for dataset composed of 12 cancers. (**a**) F-score for cDriver (solid blue line) and four other driver identification methods using the Pancan12 dataset (**b**) F-score for an ensemble approach of all tools with and without our Bayesian model, cDriver (blue and green lines respectively). (**c**) F-score for cDriver using: (i) only a published background model, (ii) including functional impact (FI), (iii) including cancer cell fraction, CCF, and (iv) a combination of all signals. (**d**) We ordered the top 30 cDriver-ranked genes on Pancan12 by their median CCF. (**d**) Matrix showing whether these top 30 genes were predicted as significant by the other four algorithms (Q value or FDR less than 0.1).

To assess whether cDriver contributes to combinations of complementary methods^28^, we calculated two ensemble F-score curves using all methods with and without cDriver (**Online methods**). Inclusion of cDriver increased the F-scores especially in the long tail between ranks 150 and 800 (Fig. 4b). Likewise, to evaluate the contribution of CCF integration to the performance of our method, we benchmarked it using only recurrence, recurrence and functional impact, recurrence and CCF, and the combination of all three signals (**Online methods**). We observed that CCF and functional impact independently improve F-score but show best performance in combination, corroborating the importance of cancer cell fraction for the identification of mutational drivers (Fig. 4c).

The top 30 genes predicted by cDriver showed a high median CCF, although with a large variance (Fig. 4d). Despite all of these genes were present in the gold standard, several of them are missed by other methods (**Fig 4e**). For example, oncodriveFM missed *FLT3* due to a cluster of medium impact mutations (**Supplementary Fig. 5a**) and OncodriveCLUST missed several tumor suppressor genes, since loss of function mutations in these genes do not necessarily cluster (e.g. *PBRM1*, **Supplementary Fig. 5b**). MutsigCV missed *KEAP1* and *MTOR* in this dataset, but it was able to find these genes with larger sample size^13^.

We next investigated whether cDriver identifies a particular function commonly missed by oncodriveFM by looking at significant genes in only one of the methods using Gene Ontology analysis (**Supplementary Fig. 12a**). Interestingly, we found that tyrosine kinase related genes were enriched in the group of genes predicted only by cDriver **(Supplementary Fig. 12c)**. Tyrosine kinase related genes have a well-described role in tumor initiation, mostly associated to oncogenes^61^, reflecting the importance of CCF as an independent discriminatory signature in addition to functional impact (oncodriveFM). Conversely, processes enriched in the group of genes predicted only by oncodriveFM were related to binding activity (**Supplementary Fig. 12d)**.

In summary, cDriver performed favorably in individual tumor types and in Pancan12 independently of applied GS, allowing us to explore an extended landscape of driver genes across multiple tumor types.

### Tumor type – driver gene landscape across 21 cancers

To obtain a comprehensive list of driver genes in an extended TCGA dataset, we ran cDriver on each and on a pooled set of 21 tumor types comprised of 6,870 samples (Pancan21, **Supplementary Table 4**). While on average the median number of significant driver genes detected per tumor type was 31 (**Supplementary Table 6**), we observed that several genes in the ‘long tail’ were a) close to significance, b) mutated in a large fraction of patients, and c) previously reported as cancer drivers in other tumor types. We therefore hypothesized that well-known driver genes with a strong positive selection signature in one or more tumor types have been missed in other cancers due to a weaker selection signature. To test this hypothesis we created a list of 216 genes with a strong positive selection signature, i.e. genes that were significant (FDR<0.1), highly ranked (top 10 of at least one tumor type or top 200 in Pancan21) and penetrant (at least 2% affected patients in at least one tumor type or across Pancan21). We next screened for these genes in the long tail of each cancer type (up to rank 100), ultimately defining a “tumor type-driver gene” (TTDG) landscape composed of 511 TTDG connections (**Supplementary Table 5, Supplementary Figure 7**).

We investigated whether the TTDG landscape reveals novel connections between driver genes and tumor types not previously published. First, we assessed the number of PubMed records obtained when querying each of the 216 genes using the MeSH term “*neoplasm*”, resulting in 189 (87%) genes with at least one and 141 (65%) genes with at least five *neoplasm*-related PubMed records. Next, we queried the gene name in combination with each specific tumor term (**Online methods**). We identified 98 (20%) novel TTDG connections consisting of 63 genes with no publication reporting recurrent somatic mutations in the connected tumor type (**Supplementary Table 7**). Furthermore, the network of these genes had significantly more interactions than expected in the STRING database (Adj. P value=2.22e-15, **Supplementary Fig. 8**). Surprisingly, 18 of these interacting genes were annotated as chromatin modifiers in the gene ontology database (Adj. P value=1.3e-10, STRING) and were significantly enriched also when considering only cancer genes (18 out of 63 versus 78 out of 504 CGC, Fisher’s exact P Value=0.0125), revealing an underappreciated role of chromatin modification and chromatin organization in several tumor types. Importantly, we found that in 18 out of 21 tumor types, 20% to 80% of patients were affected by a mutation in one of these chromatin-modifying proteins (**Supplementary Fig. 9**), with an average of 40.2% across all tumor types.

Finally, we individually investigated the TTDG connections of two known therapeutic targets, *CHD4* (Fig. 5a) and *SMARCA4* (Fig. 5b). We found that chromodomain helicase 4, *CHD4*, acts as a driver for seven tumor types, while initially it was only associated to endometrial^41^ and ovarian carcinoma (Fig. 5c). *CHD4* is a tumor suppressor and core member of the nucleosome remodeling and deacetylase (NuRD) complex^42^, which has been linked to multiple cellular processes including cell cycle regulation, DNA damage repair, and chromatin stability^43,44,45^. Survival analysis showed that bladder carcinoma patients with mutations in *CHD4* have better prognosis than patients with other mutated drivers (Fig. 5e). *SMARCA4* has a known role in lung cancer and esophageal carcinoma, and it was found recurrently mutated in pancreatic, breast, lung, and prostate cancer cell lines^46^. However, the importance of this gene as a driver in tumorigenesis has been neglected in others cancers such as head and neck and liver carcinomas (Fig. 5d). It is the core subunit of a SWI/SNF complex and has several binding motifs to other tumor suppressors proteins^47,48^. Most of the mutations fall in the active domains *SNF2* (Fig. 5b) involved in the unwinding of the DNA. Additionally, we observed that liver carcinoma patients carrying a mutation in *SMARCA4* have a poor prognosis (Fig. 5f). In summary, all novel TTDG connections could be exploited as potentially therapeutic targets ultimately increasing the number of options for cancer prognosis.

**Figure 5.**
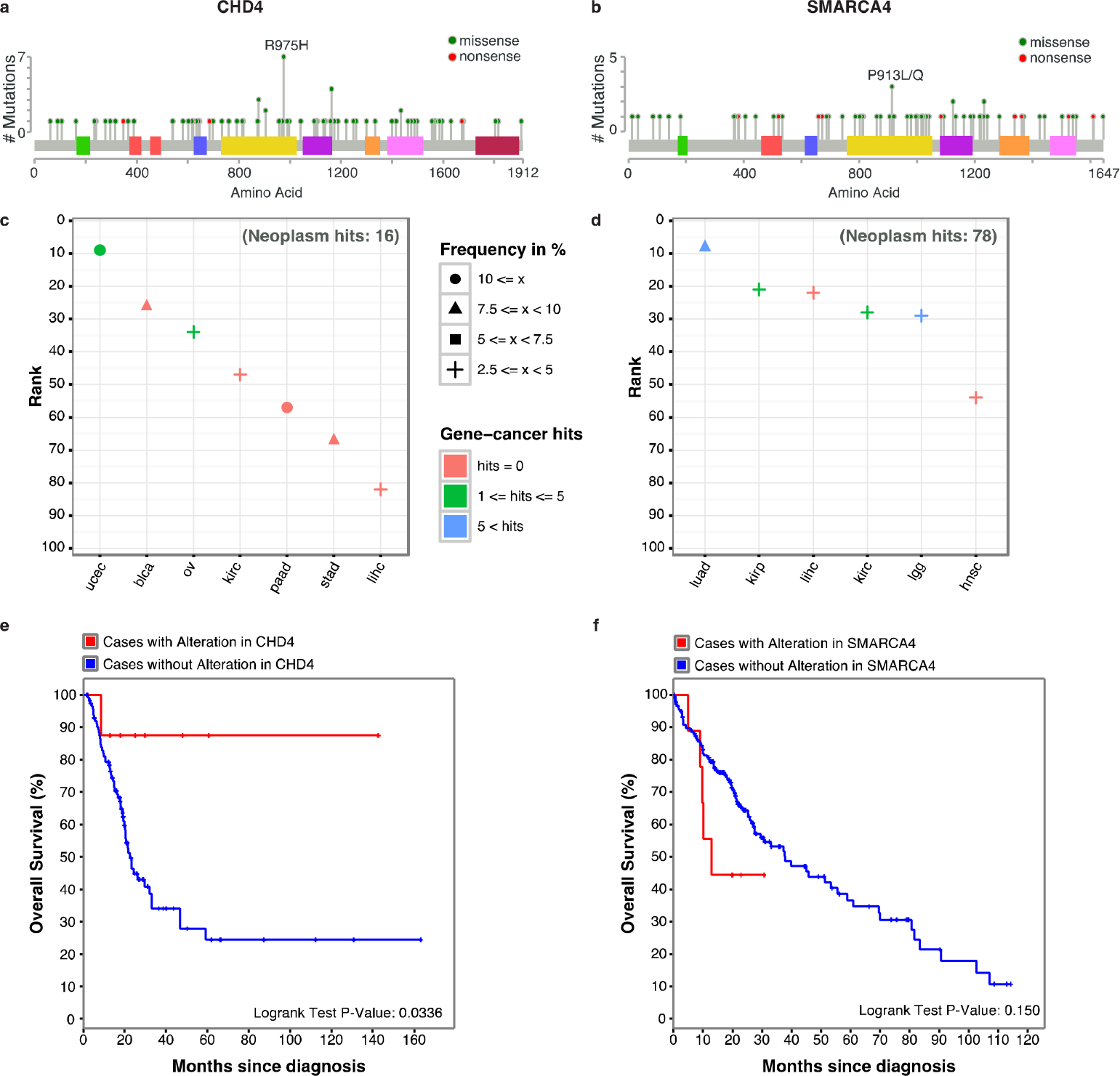
Novel *tumor type - driver gene* (TTDG) connections, CHD4 and SMARCA4. Distribution of somatic mutations found in **(a)** *CHD4* and **(b)** *SMARCA4*. The domains are colored following the cBioPortal color scheme. Most of the mutations are evenly distributed in *CHD4*, except for two small clusters at the beginning of the protein. In the case of *SMARCA4*, mutations tend to accumulate in the domains for ATP hydrolysis or DNA unwinding. TTDG connection landscape for (c) *CHD4* and (d) *SMARCA4*: the color indicates the number of pubmed hits related to each MeSH term. The shape indicates the frequency of patients affected by a mutation in the gene. Survival curves for (e) *CHD4* in bladder carcinoma and for (f) *SMARCA4* in liver hepatocellular carcinoma. Patients affected by a mutation are plotted in red.

## Discussion

Evolutionary signatures are imprinted in tumor genomes, and cDriver leverages them at the population, cellular, and molecular level to identify cancer driver genes. For the first time, we integrate these measures into a Bayesian framework to detect driver genes in three different tumor datasets, and to discover 98 unreported “tumor type - driver gene” connections across 21 tumor types. We show that these novel connections are strongly enriched for chromatin modifying proteins and have prognostic relevance, revealing an unexplored landscape of therapeutic targets.

Tumor heterogeneity complicates the discovery of cancer driver genes. Somatic mutations in these genes are under different selective pressures leading to complex tissue- and patient-specific clonal structures. Previous studies have shown that cohort recurrence and functional impact represent evidence of positive selection at the population and molecular level, respectively^49,28^. On one hand, the number of patients carrying a mutation in a gene hints at the importance of this gene for cancer etiology. On the other hand, mutations severely affecting protein function are more likely to be relevant for tumor formation^50^. Remarkably, while former studies have demonstrated that driver mutations rise in frequency within the tumor cell population^51,25^, while passenger mutations accumulate neutrally following a powerlaw distribution^52^, cancer cell fraction of mutations has been neglected as a feature for cancer driver prediction.

We found that integrating cancer cell fraction in our model increases the number of true driver genes detected. This makes an intuitive sense because positively selected (driver) mutations are expected to be present in a large fraction of cancer cells. While this can also be the case for hitchhiking passenger mutations, we demonstrate that nonsilent driver mutations are found at a significantly higher CCF than all other types of mutations. The difference in CCF indicates that most passenger mutations occur after the driver mutations and accumulate in a neutral fashion, as previously suggested^25,52^. In summary, driver mutations accumulate as a function of selection and time, while passenger mutations accumulate solely as a function of time. On the other hand, it is possible that CCF also reduces the impact of technical artifacts arising from low allele fraction of false positive calls, but such effect should be independent of the mutation type. The accuracy of CCF calculation depends on correct estimates of tumor purity and tumor ploidy, as well as adequate coverage of mutated positions. We expect that ultra-deep and single cell sequencing will further improve the power to detect mutations in small fractions of the tumor, making CCF an indispensable feature for accurate driver gene prediction.

Different types of driver genes are functionally constrained by different evolutionary pressures. Therefore, combining complementary signatures for mutational driver identification^28^, we show that functional impact and CCF equally improve performance, and their combination outperforms the use of each independently. Interestingly, genes missed by oncodriveFM (functional impact bias) but identified by cDriver (using CCF and functional impact) are enriched in a well-known group of genes involved in tumor initiation, the tyrosine kinases. These results indicate that selection signatures at the molecular, cell, and population level are complementary, likely due to different underlying biological principles. In addition, we demonstrate that cDriver improves performance when combined with canonical methods, ultimately detecting infrequently mutated driver genes missed by other approaches.

The total number of cancer genes driving tumorigenesis is still incomplete. Multiple gold standard datasets have been assembled in the literature (from 100 to 600 genes), none constituting a definite set of cancer driver genes. Although this is a limitation when benchmarking different methods, we show that the performance order is consistent across five applied gold standards. Moreover, in this study none of the methods achieved a recall higher than 30% against Cancer Gene Census, suggesting that many genes have a role in tumorigenesis not related to positive selection of nonsilent point mutations. Indeed, all methods tested here neglect other types of complex variation that may be driving cancer malignancy. These events are also under selection such as, positive selection of copy number alterations, fusion genes, regulatory, and synonymous mutations, as well as negative selection of cancer essential genes.

Our study also highlights the importance of prior information on driver gene prediction. A gene that is highly mutated (known driver) in one tumor type is probably a driver in other tumor types, even if it is infrequently mutated. We show that 63 genes highly ranked in one tumor type have been neglected in other cancers, despite a low, but substantial number (up to 10%) of affected patients. Interestingly, this list of novel “tumor type – driver gene” connections includes 18 chromatin-modifying proteins, extending the well-known and important role of chromatin remodeling in cancer^46,44^. These genes are global regulators of transcription activity and often act in a tissue-specific manner. According to a network analysis, all 18 genes are interacting or co-localized, suggesting that a single hit is needed to drive tumorigenesis. Indeed, we found that mutations in the 18 chromatin-modifying proteins affect a large fraction of cancer patients (**Supplementary Table 7**). The genes *CHD4* and *SMARCA4* demonstrate how the landscape of tumor type – driver gene connections can be exploited to identify novel therapeutic targets, especially for patients without a canonical driver mutation.

In conclusion, we show that an extensive landscape of therapeutic targets awaits exploration. We demonstrate that integrating cellular prevalence of somatic mutations as part of multiple signatures of tumor evolution allows for improved discovery of driver genes. As a result, it facilitates identification of novel tumor type - driver gene connections, which are key for improved cancer diagnosis, monitoring, and targeted treatment selection.

## Methods

Methods and any associated references are available in the online version of the paper.

## Acknowledgements

We thank Nuria Lopez-Bigas for fruitful discussion and Michael Schroeder for running oncodrive suite algorithms in the benchmark datasets. We acknowledge support of the Spanish Ministry of Economy and Competitiveness, ‘Centro de Excelencia Severo Ochoa 2013-2017’. We acknowledge the support of the CERCA Programme / Generalitat de Catalunya. This project has received funding from the European Union’s Horizon 2020 research and innovation programme under grant agreement Nº 635290. Luis Zapata has been supported by the International PhD scholarship program of La Caixa at CRG.

## Author contributions

L.Z., H.S., S.O. and X.E. conceived and designed the project. H.S. implemented the R-package. L.Z. and H.S. performed the analysis and prepared figures. O.D. preformed CNV analysis on CLL samples. M.F. performed expression analysis on CLL data. L.Z., S.O. and H.S. wrote the manuscript with the help of all the authors.

## Online Methods

### Data

Pancan12 somatic mutation data. Filtered MAF files from 12 tumor types were obtained from synapse (syn1729383). Allele counts, ploidy status, and histological purity estimates were merged into a single MAF file containing information for 3,276 samples and 617,354 mutations described elsewhere^32^. Allele counts for 782 samples not available from synapse were obtained from DCC-Firehose MAF files. Damage probability scores were added by applying a sigmoid transformation to CADD scores^53^ with mean *μ* = 15 and scale factor of 2. Individual MAF files for each cancer were produced in order to perform downstream analysis. Expression values for this dataset were also obtained from synapse (syn1729383).

Pancan21 somatic mutation data. The MAF files for 6,485 exome samples from 20 tumor types were downloaded from DCC-Firehose and combined with 385 CLL cases to obtain a large dataset of 21 tumor types (**Supplementary Table 4**). Allele counts were transformed to VAF and CADD scores were added for each mutation. We removed duplicated samples and updated the gene symbols using the Hugo Gene Symbol database. Colon and rectal tumors were merged into one tumor type giving us a final set of 20 tumor types. All curated MAF files used in this study were uploaded to synapse (syn5593040).

CLL somatic mutation data. 385 CLL tumor-normal pairs sequenced by WES were analyzed using an in-house pipeline. Reads were aligned to hg19 using BWA-mem^54^ and BAM files were post-processed (indel realignment, base quality recalibration) using GATK (https://www.broadinstitute.org/gatk/). Mutect^55^, Indelocator (https://www.broadinstitute.org/cancer/cga/indelocator), and ClinDel (unpublished) were used to produce a set of somatic SNVs and Indels. 27,625 mutations were annotated using eDiVA (www.ediva.crg.eu) to obtain several features including effect on gene and protein sequence, allele counts, CADD functional impact score, and population allele frequencies. Somatic SNVs that have a high number of occurrences across all paired normal samples, i.e. are likely germline variants (ND occurrence > 10), or a high rate of exclusion by MuTect across all samples (more than five times excluded) were excluded from the analysis. Indels falling within 30 bp of a repeat masked region were removed. Additionally, to reduce common false positive in detecting indels, we excluded indels that were reported in exons not expressed in B-lymphocytes. We used tophat and cufflinks to calculate FPKMs for 270 CLL RNA-seq samples. We considered exons not to be expressed if they had an average or median fragment per kilobase per million (FPKM) < 1. Finally, we produced a MAF file excluding variants in segmental duplications, common in the population (AF in EVS or 1000GP > 1%) or with alternative allele fraction (VAF) < 0.05. CADD damage score and VAF were added to each mutation in the final data. Ploidy values and cancer cell fraction of CNVs were obtained using the in-house developed tool clinCNV (unpublished).

### CCF distribution analysis

We estimated cancer cell fraction of mutations for Pancan12, Pancan20, and CLL-ICGC mutation data using the function CCF of the cDriver package. For each tumor type we obtained the current set of significant driver genes from the IntoGen website, having at least one significant prediction in the pooled and the individual cancer analysis. These drivers have been predicted based on recurrence (mutSigCV), functional damage bias (oncodriveFM), or clustering (oncodriveClust), but none of the predictors considered CCF. Somatic mutations were divided into 4 sets, including nonsilent mutations in driver genes, nonsilent mutations in passenger genes, silent mutations in driver genes, and silent mutations in passenger genes. We compared the CCF distribution for each group using Wilcoxon-Mann-Whitney statistical test for each cancer type separately (based on the most recent TCGA/ICGC releases) as well as for curated (Pancan12) and non-curated (Pancan16) pan-cancer sets. In this analysis we only included tumor types for which at least 10 significant drivers were found (16 tumor types in total).

### cDriver package

We have developed cDriver, an R package to identify mutational driver genes using NGS data from cancer genome studies (**Supplementary Fig. 1**). cDriver uses a MAF file (v2.4) as input data with additional mandatory column (i) variant allele frequency (VAF), and optional columns (ii) damage score, (iii) ploidy, (iv) histological purity, and (v) cancer cell fraction of the CNV. These measures can be obtained from current cancer genome or exome sequencing studies and public genome annotation databases. cDriver can use any mutation annotated as indel, missense, nonsense, splice, TSS, nonstop, and silent variant following the MAF column variant type.

One of the conceptual advances of our method is the inclusion of cancer cell fraction (CCF). Although any current method for CCF calculation can be used with cDriver, we provide a simple function to estimate CCF of SNVs and indels based on VAF, ploidy, CCF of overlapping CNVs, and tumor purity. The CCF estimation by cDriver highly correlates with the cellular prevalence calculated by PyClone^18^, while taking only seconds to compute for the complete TCGA pan-cancer data.

To account for the variability of the background mutation rate (bmr) between genes, cDriver uses silent mutations to locally estimate the expected number of nonsilent mutations. To this end, we applied a classical formula (Ka/Ks or dNdS ratio) to detect selection bias in comparative genome analysis to incorporate CCF of silent mutations. However, somatic mutations are rare for most cancers and many genes do not harbor silent mutations, restricting the usefulness of Ka/Ks. Therefore, cDriver calculates an average bmr using the CCF-adjusted Ka/Ks formula and a pre-calculated bmr taken from the literature^13^.

Next, cDriver calculates posterior probabilities per gene using two Bayesian models, (i) the “cancer hazard model” and (ii) the “driver model”. The first model requires the incidence of the tumor in the population as prior probability, while the second requires a list of known driver genes (e.g. any gold standard used in this study) to estimate prior and likelihood values. The “cancer-hazard model” estimates the posterior probability of developing cancer if the focal gene is mutated, given evidence from the data, i.e. somatic mutations of the gene in a cohort of cancer patients. The “driver inference model” estimates the posterior probability that a gene is a true cancer driver given evidence from the data.

As a final step, cDriver can provide an optimum rank-cut off value by estimating FDR at each rank based on a null model. cDriver default estimates the optimum rank for each Bayesian model and suggest the best rank cut-off as the average value between both ranks.

In summary, cDriver combines recurrence, CCF, and functional impact as a foreground measure, and an averaged background mutation probability as a measure to calculate posterior probabilities for each gene. In the next sections each step is described in detail.

## Step 1) Estimation of cancer cell fraction per gene

### CCF calculation

We developed a function for estimation of Cancer Cell Fraction (CCF) per gene as part of the cDriver package (for details see **Supplementary note**), although any method can be used to estimate the CCF subsequently used for driver prediction (e.g. PyClone). Intuitively, we assumed that the variant allele should be observed in approximately half of the reads if it is a clonal heterozygous variant in a diploid locus. In this case, CCF is calculated as the variant allele frequency (VAF) multiplied by two and corrected for the purity of the cancer sample. All other cases are described in the supplementary note.

## Step 2) Background mutation rate models

### CCF-adjusted counts of nonsilent and silent mutations

First, we adjusted the classic formula for detecting selection from comparative data^56^ to estimate the expected number of nonsilent mutations under no selective pressures (i.e. neutral evolution). This formula is the ratio between the rate of nonsynonymous substitutions (*n_a_*) per nonsynonymous sites (*N_a_*) and the rate of synonymous substitutions (*n_s_*) per synonymous sites (*N_s_*):

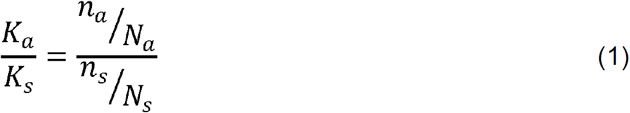

Next, we adapted the formula to take into account cancer cell fraction of mutations by calculating *n_s_* and *n_a_* as the sum of CCF of silent and nonsilent mutations, respectively, resulting in 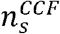 and 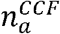. The CCF-adjusted *K_a_*/*K_s_* formula is:

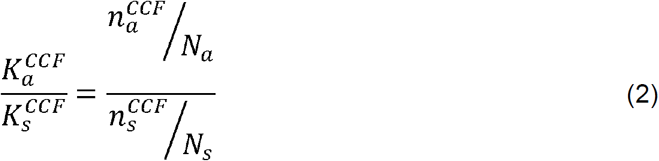

### Expected number of nonsilent mutations per gene

We estimated the expected 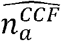 for each gene in a cancer cohort-specific manor based on the observed number of CCF-adjusted silent mutations in coding regions 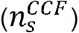 within the provided cohort (e.g. WES data from BRCA, CLL, Pancan12, Pancan21). The total number of sites (*N_a_* and *N_s_*) was taken from Lawrence et al^13^. Under the assumption of neutral selection 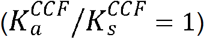, we estimated an expected 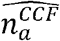 as:

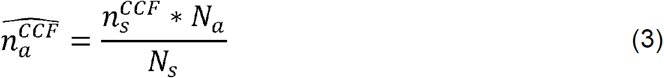

### Zero counts of silent mutations

To avoid zero or very low 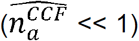 expected nonsilent mutations in equation 3, we defined a minimum 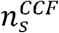. We assume that one nonsilent mutation per gene in the cohort can occur by chance and should not be considered a positive selection signal. Thus, for a gene with zero silent mutations observed (or where CCF of silent mutations is very low), we added a pseudocount to silent mutations such that equation (2) 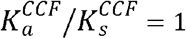 (neutral), assuming one nonsilent mutation, and recalculated 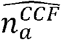 in equation (3).

### Background mutation probability based on the expected number of nonsilent mutations

After obtaining the expected 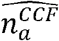 for every gene we calculated the probability that a patient has at least one nonsilent mutation in a gene X, P(X>=1). To this end, we approximated the average number of somatic nonsilent mutations in a healthy cohort (*r*) using the cancer cohort. Here, we assume that the majority of clonal mutations detected (CCF_SNV_ ≥ 0.85) are passengers present before tumor initiation. We recognize that the number of somatic nonsilent mutations not caused by cancer varies from patient to patient. We set the number of trials, r, to the average number of mutations per patient in the cohort. In one trial the probability of success is given by 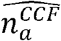 for gene X divided by the sum of 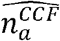 across all genes. Following this assumption, we estimated the probability P(*X* ≥ 1) in r trials using a binomial distribution:

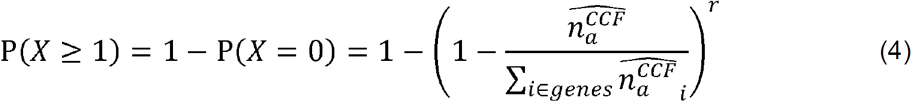
where P(*X* = 0) is a probability for a gene X to have no nonsilent mutations, and derived as 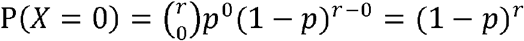 where p is probability of success 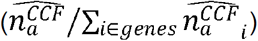.

### Background mutation probability based on other mutation rate estimates

To compensate for the lack of power of the CCF-adjusted *K_a_*/*K_s_* model for genes, we additionally incorporated the non-coding mutation rate (*ncmr*) provided by Lawrence et al^13^. For each gene the *ncmr* and the gene length were used to calculate the number of total expected mutations (silent and nonsilent) 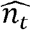. Under the assumption of neutral evolution (*K_a_*/*K_s_* = 1), we determined the expected number of nonsilent mutations following a rearrangement of the classical formula into:

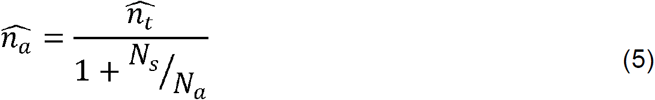

Similarly to the previous model, we calculated the probability that a gene has at least one nonsilent mutation using the binomial distribution formula, but without CCF:

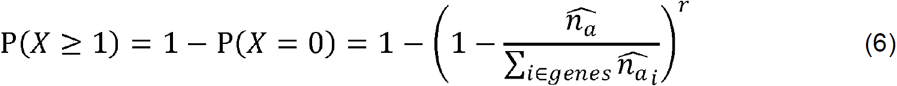

Given the continuous progress on technologies, it is likely that more specific somatic mutation rates will be calculated in the future, so in addition cDriver could integrate any measure of background mutation rate. Here, we used as final background mutation probability (bmp) the average bmp obtained by the two methods described above (CCF-adjusted *K_a_*/*K_s_* and *ncmr*).

## Step 3) Bayesian inference models

### Cancer-hazard inference model

In the first model, we adapted Bayes formula to calculate a posterior probability of developing cancer given that a focal gene is mutated as

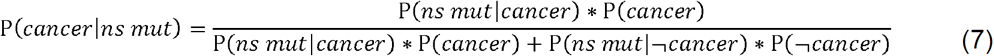
where the prior probability for developing cancer, P(*cancer*), is the incidence of the cancer type in the population. The likelihood, P(*ns mut|cancer*), that a cancer patient carries a nonsilent mutation in a gene of interest is estimated from the cohort. To this end, we used the sum of CCF times the adjusted-CADD damage probability per gene across all patients:

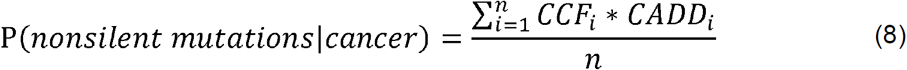
where *i* is the index of the patient and *n* the total size of the cohort. If a patient did not have any nonsilent mutation, then CCF was equal to zero. If two nonsilent mutations were found in a patient in the same gene of interest, we used the mutation with the highest CCF.

We defined the marginal probability of having a nonsilent mutation as the sum of the numerator of (1), plus the conditional probability of having a nonsilent mutation in a healthy population P(*ns mut|¬cancer*) times the probability of a healthy individual P(¬*cancer*). We denoted P(*ns mut|¬cancer*) as the somatic background mutation probability (bmp). To our knowledge there is no large enough cohort of healthy people examined for tissue specific somatic mutations, therefore direct estimation from data is not possible. However, we estimated an upper bound of the bmp as described in the previous section.

### Driver inference model

In the second model, we calculated the posterior probability that a gene is a cancer driver given the mutation data in the studied cohort using the formula:

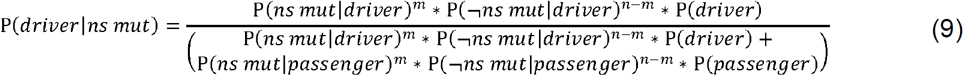
where *m* is equal to 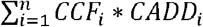 and *n* is total size of the cohort.

To estimate a prior probability of a gene being a driver we need to consider that most tumor types can be caused by mutations in a different set of genes. Depending on which tumor type or group of cancers (‘pan-cancer’) we were analyzing, the number of known driver genes differs and hence the prior probabilities change (e.g. ovarian cancers are in most cases caused by mutation in TP53, while the number of published genes involved in CLL ranges from 20 to 40, depending on the study). We estimated the prior probability that a random gene is a driver as equal to the ratio between the number of known driver genes of the cancer type and the total number of protein coding genes:

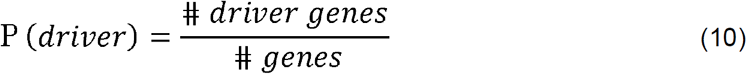

The number of driver genes can be approximated as the number of published driver genes for a particular cancer type. If the cancer has not been studied yet, or if we deal with pan-cancer sets of multiple cancer types, the prior can be approximated using any gold standard list of cancer driver genes.

Because of inter-tumor heterogeneity genes that are known to be cancer drivers in a given tumor type will not necessarily be mutated in all patients. The probability that a gene is mutated given that it is a known driver can be estimated as:

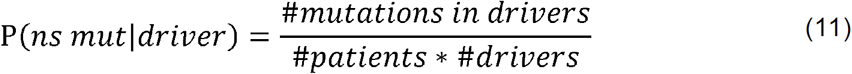
where we assume that all drivers have the same chance to be mutated. As this assumption is weak the cDriver package allows the user to define better estimates for this likelihood.

The probability that a gene is mutated given that it is not a driver P(*ns mut|passenger*) is estimated from the background mutation rate as described previously.

The other terms in the equation, P(¬*ns mut|driver*) and P(¬*ns mut|passenger*) were calculated as the complementary events of P(*ns mut|driver*) and P(*ns mut|passenger*) described above.

## Step 4) Optimum rank cut off selection

### Significant rank selection based on weighted sampling of a null model

To generate an optimum rank cut off for our two bayesian models, we calculated a null model based on random assignment of new gene labels based on the background mutation probability (bmp) vector. The BMP vector is generated from the observed silent mutation data as described previously. Then, we run our Bayesian model as we would with the cancer cohort to obtain posterior probabilities per gene under a null model. We repeat this 100 times to be sure that probabilities are stable between each run and we are not catching unlucky random assignment of gene names. Finally, cDriver calculates the optimum threshold by comparing the ranking of the true set versus the null model, assuming a false discovery rate of 10% (FDR < 10%, any value can be selected by the user)

### Running competing methods for cancer driver gene identification

(i) MuSiC on Pancan12 and BRCA: gene coverage files and results from the MuSiC suite for the Pancan12 dataset (including all BRCA cases used here) were obtained from synapse (syn819550, syn1713813, syn1734155). For this set of results, we only considered genes that were less than 0.05 FDR in at least two out of three measures. The rank order was based on the P-values given by the CT test, followed by the LRT test, followed by the number of cases affected. MuSiC on CLL dataset: gene coverage files for 385 samples were generated from tumor-normal BAM files using the function calc-bmr available in the MuSiC suite. The region of interest files (ROI) were downloaded from synapse and merged to avoid duplicates. For comparative analysis we used the sorted list returned by the tool. (ii) MutSigCV on all datasets: we ran MutSigCV using default parameters on all datasets, assuming full coverage and using the example covariate space provided in the source code. (iii) OncodriveClust and (iv) oncodriveFM: The analysis was performed by the group of Nuria Lopez-Bigas. (v) cDriver on all dataset: We ran cDriver using default parameters. Prior values used for the cancer-hazard model are shown in Supplementary Table 1 and 3.

### Benchmarking

For comparative analysis of several competing methods we tested datasets that differed in the number of samples, the number of mutations, the tissue-of-origin, and the purity of the tumor on different gold standard datasets. We downloaded published lists of significantly mutated genes (SMGs) from: 1) Kandoth et al., 2013^32^, 127 significantly mutated genes across 12 tumor types; 2) Lawrence et al., 2014^30^, 261 cancer genes predicted from 21 tumor types; 3) Tamborero et al., 2013^28^, 435 cancer genes predicted by a combination of multiple algorithms. For benchmarking purposes, we only used the 291 genes labeled ‘high confident’; 4) Cancer Gene Census^39^, list of 547 manually curated cancer driver genes; 5) Xie et al., 2014^40^, list of 556 cancer-associated genes; 6) Landau et al., 2013^8^, a list of 21 CLL specific genes and 7) TCGA breast 2012^57^, 35 breast cancer genes.

The gold standard genes for breast cancer consisted of the union of dataset (7), and breast cancer genes found in (2) and (4) plus the top 20 genes identified in COSMIC. For CLL we merged dataset (6) and CLL genes found in (2) and (4) plus the top 20 genes identified in COSMIC. Subsequently, we manually curated this list by checking the number of PubMed records found by querying the HUGO gene name and the corresponding MeSH term for breast cancer and CLL. We excluded histone genes that have no relevant publication associating them to recurrent somatic mutations. Results for Pancan12 were benchmarked using datasets 1-5. Single tumor types (i.e. CLL and breast cancer) were benchmarked against the tumor specific gold standards assembled as described above. In addition, we compared the performance of the methods on Pancan12 with and without filtration of non-expressed genes.

Furthermore, we benchmarked our method cDriver under several scenarios and parameter settings: the F-score curve for (i) cDriver using a simple recurrence model where no CCF or functional impact was used and the background model did not include CCF-adjusted Ka/Ks, (ii) cDriver using only functional impact, (iii) cDriver using CCF adjusted mutation counts and the CCF-adjusted Ka/Ks background model, and (iv) cDriver using all signatures of positive selection. Lastly, we benchmarked an ensemble of complementary methods (MuSic, MutSigCV, OncodriveFM, OncodriveClust) including and not including cDriver. For this, we calculated a combined rank and calculate the Fscore. We used “*Borda count”* ranking method with truncated ranks (up to two times the gold standard size) but using ranks of only the three best methods.

For visualization purposes we show only genes that were ranked up to twice the number of gold standard genes given no further improvement was achieved by any tool beyond these thresholds.

### Defining the landscape of tumor type–driver gene connections in Pancan21

We ran cDriver on each of the 21 tumor types separately (syn5593040, **Supplementary Table 4**) and in the pooled Pancan21 dataset. Next, we used the union of the top 10 genes for each tumor type (except for ovarian and thyroid carcinoma) and the top 200 genes of the pooled Pancan21 cDriver results to create a list of high confidence driver genes. For each of these genes, we noted their presence among the top 100 ranked genes of each tumor type to define a “tumor type – driver gene” (TTDG) connection. We defined genes found in the top 10 of only one tumor type and not in the top 100 of any other tumor type as highly tumor specific.

### Identification of novel TTDG connections by PubMed mining

For each high confidence gene, we queried the HUGO symbol together with the MeSH term “neoplasm” (i.e. “ATM[TIAB] AND neoplasm[MH]”) against the PubMed database in order to test if the gene have been associated to any cancer type. The HUGO symbol had to be found in the title and/or abstract. Next, for each TTDG connection we queried the HUGO symbol together with the MeSH term of the associated tumor type. Based on the PubMed mining results, we used the following criteria to detect novel TTDG connections: (i) the tested gene was among the top 200 of the Pancan21 analysis, (ii) the gene had at least 5 *neoplasms*-related PubMed records, (iii) the gene was among the top 100 of the corresponding tumor type (defined by its TTDG connection), (iv) the gene had zero or one TTDG specific PubMed record, and, (v) if one TTDG specific PubMed entry was retrieved, we required that the publication did not report recurrent somatic mutations for that gene in the tumor type of interest.

### Protein interaction and functional enrichment analysis

We used STRING v10.0^58^ to find the connectivity among the novel TTDGs reported. We input the list of genes into the webserver and retrieved the network using all STRING features except text-mining and database evidence. STRING provides built-in analysis functions to detect protein-protein interactions and to perform GO term enrichment analysis. For the latter, STRING performs a Hypergeometric test and corrects for multiple testing using Benjamini and Hochberg. GO term enriched in the analysis of the specific oncodriveFM results only versus cDriver results only were input into the analysis platform REVIGO ^59^ to find semantically relevant terms.

### Individual gene analysis

To visualize the somatic mutations on the gene structure we used MutationMapper^60^. We input our list of nonsilent mutations for genes individual genes from the Pancan12 dataset. The clinical data (defined as processed data, level 2, by TCGA) for the patients was downloaded from the TCGA data portal (https://tcga-data.nci.nih.gov/tcga/). Kaplan-Meier curves and log-rank p-values for the selected genes were calculated using the R package Surv, patient-gene pairs were classified as affected if they presented at least one of these types of mutations ‘Frame_Shift_Del’, ‘Frame_Shift_Ins’, ‘In_Frame_Del’, ‘In_Frame_Ins’, ‘Missense_Mutation’, ‘Nonsense_Mutation’, ‘Splice_Site’, ‘Translation_Start_Site’, ‘Nonstop_Mutation’. Multiple testing correction was performed using Benjamini and Hochberg for the selected connections of chromatin modifiers.

### Code availability

The latest version of the cDriver R package, with documentation and example data sets, is freely available at github.com/hanasusak/cDriver.

## References

1. Nowell, P. C. The clonal evolution of tumor cell populations. Science 194, 23–28 (1976).

2. Stratton, M. R., Campbell, P. J. & Futreal, P. A. The cancer genome. Nature 458, 719–724 (2009).

3. Vogelstein, B., Papadopoulos, N., Velculescu, V. E., Zhou, S., et al. Cancer genome landscapes. science 339, 1546–1558 (2013).

4. Fearon, E. R. & Vogelstein, B. A genetic model for colorectal tumorigenesis. Cell 61, 759–767 (1990).

5. Sakoparnig, T., Fried, P. & Beerenwinkel, N. Identification of constrained cancer driver genes based on mutation timing. PLoS Comput Biol 11, e1004027 (2015).

6. Bissell, M. J. & Hines, W. C. Why don’t we get more cancer? A proposed role of the microenvironment in restraining cancer progression. Nat Med 17, 320–329 (2011).

7. Beerenwinkel, N., Schwarz, R. F., Gerstung, M. & Markowetz, F. Cancer evolution: mathematical models and computational inference. Systematic biology 64, e1–e25 (2015).

8. Landau, D. A., Carter, S. L., Stojanov, P., McKenna, A., et al. Evolution and impact of subclonal mutations in chronic lymphocytic leukemia. Cell 152, 714–726 (2013).

9. Gerlinger, M., Rowan, A. J., Horswell, S., Larkin, J., et al. Intratumor heterogeneity and branched evolution revealed by multiregion sequencing. New England Journal of Medicine 366, 883–892 (2012).

10. Bolli, N., Avet-Loiseau, H., Wedge, D. C., Van Loo, P., et al. Heterogeneity of genomic evolution and mutational profiles in multiple myeloma. Nat Commun 5, 2997 (2014).

11. Zhao, B., Hemann, M. T. & Lauffenburger, D. A. Intratumor heterogeneity alters most effective drugs in designed combinations. Proc Natl Acad Sci U S A 111, 10773–10778 (2014).

12. Marusyk, A., Almendro, V. & Polyak, K. Intra-tumour heterogeneity: a looking glass for cancer? Nat Rev Cancer 12, 323–334 (2012).

13. Lawrence, M. S., Stojanov, P., Polak, P., Kryukov, G. V., et al. Mutational heterogeneity in cancer and the search for new cancer-associated genes. Nature (2013).

14. Sottoriva, A., Spiteri, I., Piccirillo, S. G., Touloumis, A., et al. Intratumor heterogeneity in human glioblastoma reflects cancer evolutionary dynamics. Proceedings of the National Academy of Sciences 110, 4009–4014 (2013).

15. Lee, J.-Y., Yoon, J.-K., Kim, B., Kim, S., et al. Tumor evolution and intratumor heterogeneity of an epithelial ovarian cancer investigated using next-generation sequencing. BMC Cancer 15, (2015).

16. Oesper, L., Mahmoody, A. & Raphael, B. J. THetA: Inferring intra-tumor heterogeneity from high-throughput DNA sequencing data. Genome Biology 14, R80 (2013).

17. Fischer, A., Vázquez-García, I., Illingworth, C. J. & Mustonen, V. High-Definition Reconstruction of Clonal Composition in Cancer. Cell Rep (2014).

18. Roth, A., Khattra, J., Yap, D., Wan, A., et al. PyClone: statistical inference of clonal population structure in cancer. Nature methods (2014).

19. Miller, C. A., White, B. S., Dees, N. D., Griffith, M., et al. SciClone: inferring clonal architecture and tracking the spatial and temporal patterns of tumor evolution. PLoS Comput Biol 10, e1003665 (2014).

20. Li, S. C., Tachiki, L. M., Kabeer, M. H., Dethlefs, B. A., et al. Cancer genomic research at the crossroads: realizing the changing genetic landscape as intratumoral spatial and temporal heterogeneity becomes a confounding factor. Cancer Cell Int 14, 115 (2014).

21. Campbell, P. J., Yachida, S., Mudie, L. J., Stephens, P. J., et al. The patterns and dynamics of genomic instability in metastatic pancreatic cancer. Nature 467, 1109–1113 (2010).

22. Gerstung, M., Beisel, C., Rechsteiner, M., Wild, P., et al. Reliable detection of subclonal single-nucleotide variants in tumour cell populations. Nat Commun 3, 811 (2012).

23. Schuh, A., Becq, J., Humphray, S., Alexa, A., et al. Monitoring chronic lymphocytic leukemia progression by whole genome sequencing reveals heterogeneous clonal evolution patterns. Blood 120, 4191–4196 (2012).

24. Bassaganyas, L., Beà, S., Escaramís, G., Tornador, C., et al. Sporadic and reversible chromothripsis in chronic lymphocytic leukemia revealed by longitudinal genomic analysis. Leukemia (2013).

25. McGranahan, N., Favero, F., de Bruin, E. C., Birkbak, N. J., et al. Clonal status of actionable driver events and the timing of mutational processes in cancer evolution. Sci Transl Med 7, 283ra54 (2015).

26. Dees, N. D., Zhang, Q., Kandoth, C., Wendl, M. C., et al. MuSiC: identifying mutational significance in cancer genomes. Genome Res 22, 1589–1598 (2012).

27. Gonzalez-Perez, A. & Lopez-Bigas, N. Functional impact bias reveals cancer drivers. Nucleic Acids Res 40, e169 (2012).

28. Tamborero, D., Gonzalez-Perez, A. & Lopez-Bigas, N. OncodriveCLUST: exploiting the positional clustering of somatic mutations to identify cancer genes. Bioinformatics 29, 2238–2244 (2013).

29. Tamborero, D., Gonzalez-Perez, A., Perez-Llamas, C., Deu-Pons, J., et al. Comprehensive identification of mutational cancer driver genes across 12 tumor types. Sci Rep 3, 2650 (2013).

30. Lawrence, M. S., Stojanov, P., Mermel, C. H., Robinson, J. T., et al. Discovery and saturation analysis of cancer genes across 21 tumour types. Nature 505, 495–501 (2014).

31. Smith, J. M. & Haigh, J. The hitch-hiking effect of a favourable gene. Genet Res 89, 391–403 (2007).

32. Kandoth, C., McLellan, M. D., Vandin, F., Ye, K., et al. Mutational landscape and significance across 12 major cancer types. Nature 502, 333–339 (2013).

33. Puente, X. S., Beà, S., Valdés-Mas, R., Villamor, N., et al. Non-coding recurrent mutations in chronic lymphocytic leukaemia. Nature 526, 519–524 (2015).

34. Dieci, M. V., Smutná, V., Scott, V., Yin, G., et al. Whole exome sequencing of rare aggressive breast cancer histologies. Breast Cancer Res Treat 156, 21–32 (2016).

35. Álvarez-Silva, M. C., Yepes, S., Torres, M. M. & Barrios, A. F. Proteins interaction network and modeling of IGVH mutational status in chronic lymphocytic leukemia. Theor Biol Med Model 12, 12 (2015).

36. Nik-Zainal, S., Davies, H., Staaf, J., Ramakrishna, M., et al. Landscape of somatic mutations in 560 breast cancer whole-genome sequences. Nature (2016).

37. Landau, D. A., Tausch, E., Taylor-Weiner, A. N., Stewart, C., et al. Mutations driving CLL and their evolution in progression and relapse. Nature 526, 525–530 (2015).

38. de Miranda, N. F., Georgiou, K., Chen, L., Wu, C., et al. Exome sequencing reveals novel mutation targets in diffuse large B-cell lymphomas derived from Chinese patients. Blood 124, 2544–2553 (2014).

39. Futreal, P. A., Coin, L., Marshall, M., Down, T., et al. A census of human cancer genes. Nat Rev Cancer 4, 177–183 (2004).

40. Xie, M., Lu, C., Wang, J., McLellan, M. D., et al. Age-related mutations associated with clonal hematopoietic expansion and malignancies. Nat Med 20, 1472–1478 (2014).

41. Le Gallo, M., O’Hara, A. J., Rudd, M. L., Urick, M. E., et al. Exome sequencing of serous endometrial tumors identifies recurrent somatic mutations in chromatin-remodeling and ubiquitin ligase complex genes. Nat Genet 44, 1310–1315 (2012).

42. Cai, Y., Geutjes, E. J., De Lint, K., Roepman, P., et al. The NuRD complex cooperates with DNMTs to maintain silencing of key colorectal tumor suppressor genes. Oncogene 33, 2157–2168 (2014).

43. Lai, A. Y. & Wade, P. A. Cancer biology and NuRD: a multifaceted chromatin remodelling complex. Nat Rev Cancer 11, 588–596 (2011).

44. O’Shaughnessy, A. & Hendrich, B. CHD4 in the DNA-damage response and cell cycle progression: not so NuRDy now. Biochem Soc Trans 41, 777–782 (2013).

45. Chudnovsky, Y., Kim, D., Zheng, S., Whyte, W. A., et al. ZFHX4 interacts with the NuRD core member CHD4 and regulates the glioblastoma tumor-initiating cell state. Cell Rep 6, 313–324 (2014).

46. Roberts, C. W. & Orkin, S. H. The SWI/SNF complex‐‐chromatin and cancer. Nat Rev Cancer 4, 133–142 (2004).

47. Orvis, T., Hepperla, A., Walter, V., Song, S., et al. BRG1/SMARCA4 inactivation promotes non-small cell lung cancer aggressiveness by altering chromatin organization. Cancer Res 74, 6486–6498 (2014).

48. Biegel, J. A., Busse, T. M. & Weissman, B. E. SWI/SNF chromatin remodeling complexes and cancer. Am J Med Genet C Semin Med Genet 166C, 350–366 (2014).

49. Babenko, V. N., Basu, M. K., Kondrashov, F. A., Rogozin, I. B. & Koonin, E. V. Signs of positive selection of somatic mutations in human cancers detected by EST sequence analysis. BMC Cancer 6, 36 (2006).

50. Ostrow, S. L., Barshir, R., DeGregori, J., Yeger-Lotem, E. & Hershberg, R. Cancer evolution is associated with pervasive positive selection on globally expressed genes. PLoS Genet 10, e1004239 (2014).

51. Schuh, A., Becq, J., Humphray, S., Alexa, A., et al. Monitoring chronic lymphocytic leukemia progression by whole genome sequencing reveals heterogeneous clonal evolution patterns. Blood 120, 4191–4196 (2012).

52. Williams, M. J., Werner, B., Barnes, C. P., Graham, T. A. & Sottoriva, A. Identification of neutral tumor evolution across cancer types. Nature genetics (2016).

53. Kircher, M., Witten, D. M., Jain, P., O’Roak, B. J., et al. A general framework for estimating the relative pathogenicity of human genetic variants. Nat Genet 46, 310–315 (2014).

54. Li, H. & Durbin, R. Fast and accurate short read alignment with Burrows‐‐Wheeler transform. Bioinformatics 25, 1754–1760 (2009).

55. Cibulskis, K., Lawrence, M. S., Carter, S. L., Sivachenko, A., et al. Sensitive detection of somatic point mutations in impure and heterogeneous cancer samples. Nat Biotechnol (2013).

56. Nielsen, R. Molecular signatures of natural selection. Annu Rev Genet 39, 197–218 (2005).

57. Cancer Genome Atlas Network Comprehensive molecular portraits of human breast tumours. Nature 490, 61–70 (2012).

58. Szklarczyk, D., Morris, J. H., Cook, H., Kuhn, M., et al. The STRING database in 2017: quality-controlled protein–protein association networks, made broadly accessible. Nucleic Acids Research gkw937 (2016).

59. Supek, F., Bo\vsnjak, M., \vSkunca, N. & \vSmuc, T. REVIGO summarizes and visualizes long lists of gene ontology terms. PloS one 6, e21800 (2011).

60. Vohra, S. & Biggin, P. C. Mutationmapper: a tool to aid the mapping of protein mutation data. PLoS One 8, e71711 (2013).

